# Low replication stress leads to specific replication timing advances associated to chromatin remodelling in cancer cells

**DOI:** 10.1101/2020.08.19.256883

**Authors:** Lilas Courtot, Elodie Bournique, Chrystelle Maric, Laure Guitton-Sert, Miguel Madrid-Mencía, Vera Pancaldi, Jean-Charles Cadoret, Jean-Sébastien Hoffmann, Valérie Bergoglio

## Abstract

DNA replication is well orchestrated in mammalian cells through a tight regulation of the temporal order of replication origin activation, named the replication timing, a robust and conserved process in each cell type. Upon low replication stress, the slowing of replication forks induces delayed replication of fragile regions leading to genetic instability. The impact of low replication stress on the replication timing in different cellular backgrounds has not been explored yet. Here we analysed the whole genome replication timing in a panel of 6 human cell lines under low replication stress. We first demonstrated that cancer cells were more impacted than non-tumour cells. Strikingly, we unveiled an enrichment of specific replication domains undergoing a switch from late to early replication in some cancer cells. We found that advances in replication timing correlate with heterochromatin regions poorly sensitive to DNA damage signalling while being subject to an increase of chromatin accessibility. Finally, our data indicate that, following release from replication stress conditions, replication timing advances can be inherited by the next cellular generation, suggesting a new mechanism by which cancer cells would adapt to cellular or environmental stress.

## INTRODUCTION

DNA replication is a highly complex process that ensures the accurate duplication of the genome, hence the faithful transmission of genetic material to the cell progeny. DNA replication occurs during S phase through the replisome activity, but it requires important upstream regulation during the G1 phase and checkpoints in G2 phase, together with a tight control throughout the process itself. Multicomplex replication machinery performs the coordinated initiation of DNA synthesis at hundreds of replication origins spread throughout the whole length of the genome (1). Adjacent origins that initiate DNA replication at the same time have been called “replicon clusters” (2), giving rise to chromosomal domains replicating synchronously. Each replicon cluster starts replicating at a precise moment during the S-phase, either at the beginning (Early-S), the middle (Mid-S) or the end (Late-S). This coordination of the temporal program of DNA replication is called “replication timing” (RT), allowing a complete and faithful duplication of the entire genome before cell division.

The RT program is modified during organism development and cell differentiation (3,4) and is coupled with gene expression, chromatin epigenome and nuclear 3D compartmentalization (5–8). In somatic cells, the RT pattern is very robust through cell generations (8–11) with early-replicating DNA residing deep within the nucleus within the A compartment containing active chromatin while the later-replicating regions occur at the nuclear periphery or near the nucleolus (9, 12, 13) within the B compartment, containing inactive chromatin. Additional complex associations have been highlighted such as the link between early-replicating regions and GC nucleotides enrichment, enhanced gene expression, and active epigenetic marks corresponding to open or euchromatin. Conversely, late-replicating regions tend to be enriched in AT nucleotides, low gene content, and have heterochromatin repressive epigenetic marks (13, 14).

DNA replication stress is defined as the slowing or stalling of the replication fork resulting in inefficient DNA replication. Many exogenous or endogenous sources of impediment on DNA as well as pathological perturbations such as oncogene activation, conflicts between DNA replication and transcription or shortage of nucleotides affect the progression of replication forks, inducing replication stress (15–19). Experimentally, replication stress can be induced by the specific inhibition of replicative DNA polymerases by treatment with the drug aphidicolin. Notably, low doses of aphidicolin (0.1 to 0.6 µM) are well known to cause the induction of common fragile sites (CFS) expression and the generation of under-replicated DNA that leads to DNA damage transmission (20–24). CFS are chromosomal regions harbouring cancer-related genes (25) that are prone to breakage upon replication stress (26) and whose instability is often observed at the early stages of carcinogenesis (27). The fragility of these chromosomal regions has been widely studied, revealing incomplete DNA replication before mitosis (21, 23, 28, 29) mainly due to conflicts with large transcription units (30–32) or/and origin paucity (33, 34).

Evidence of aberrant RT in many different genetic diseases and cancers suggests this cellular process is important for genomic stability (35–37). Interestingly, replication stress inducing CFS expression also affects the RT of these specific chromosomal domains (30, 32). The extent to which these RT changes influence tumour transformation process is still largely unknown.

The major aim of this study was to explore whether low replication stress affects differentially RT of cells from diverse types, and whether a common mechanism for RT change can be found upon low RS. To do so, we characterized and compared the impact of mild replication stress induced by low doses of aphidicolin on 4 cancer and 2 non-tumour human cell lines (colon, blood, osteoblast, retina, and lung) (**Table S1**). Our experiments revealed that a low dose of aphidicolin have a stronger effect on the RT of cancer cells, promoting RT delays but also unexpected RT advances. We demonstrated that RT advanced loci can be housed in CFS but, contrary to RT delays, they are poorly targeted by DNA damage signalling while being characterized by stronger chromatin accessibility in response to aphidicolin. Finally, we observed the persistence of RT advances in daughter cells released from replication stress which is correlated with modification of chromatin loop size and pre-replication complex (pre-RC) loading in G1 and an increase in the expression of genes contained within these chromosomal regions. Altogether, our results indicate that low replication stress, which leads to RT advances onto flexible heterochromatin regions, can influence the DNA replication program and gene expression of the next generation of cancer cells.

## MATERIALS AND METHODS

### Cell lines, cell culture and drugs

The 6 human cell lines were purchased from ATCC. Cells were grown in culture medium supplemented with 10% fetal bovine serum (Gibco Life Technologies A31608-02) at 37°C, 5% CO2 and 5% O2. HCT116, U2OS and RKO cell lines were grown in Dulbecco’s Modified Eagle’s Medium (DMEM, Gibco Life Technologies 31966021), MRC5-N cell line was grown in Minimum Essential Medium Eagle (MEM-aplha, Gibco Life Technologies 22561021), RPE-1 cell line was grown in Roswell Park Memorial Institute Media (RPMI, Gibco Life Technologies 61870044) and K562 in Iscove Modified Dulbecco Media (IMDM, Gibco Life Technologies 21980032) supplemented with decomplemented serum. Aphidicolin (Sigma-Aldrich AO781-1MG) stock solution was diluted in DMSO (Sigma-Aldrich D8418-250mL) and kept at -20°C for a maximum of 2 months after first thawing. Cells were synchronized in G1/S with 0.5mM L-Mimosine (Sigma-Aldrich M0253) for 24h.

### Fluorescence Activated Cell Sorting (FACS)

Cells were pulse-labelled with 10μM BrdU and/or EdU for indicated times then collected by trypsinization and fixed in 70% ice-cold ethanol overnight at −20°C. For EdU and BrdU immunodetection, we followed the protocol described in Bradford and Clarke 2011 (38). Finally, after washing in PBS–BSA 0.5%, then in PBS, cells were resuspended in PBS with propidium iodide (25 μg/mL, Invitrogen, p3566) and RNase A (100 μg/mL, Thermo Fisher Scientific, ENO531) or with DAPI (1/1000, Sigma-Aldrich, D9542). After 20 min of incubation at room temperature, cell cycle analysis was carried out by flow cytometry with a MACSQuant 10 or VYB cytometer (Miltenyi Biotec) and analysed with FlowLogic software.

### Cell lysis, fractionation and Western blotting

For whole cell extract, cells were lysed for 30min on ice with classic lysis buffer (0.3M NaCl, 1% triton, 50mM Tris pH7.5, 5mM EDTA, 1mM DTT and 1X Halt protease and phosphatase inhibitor cocktail from Thermo Fisher Scientific 78445). For subcellular fractionation, cells were lysed in Buffer A (Hepes 10mM pH 7.9, KCl 10mM, MgCl2 1.5mM, sucrose 0.34M, Glycerol 10%, dithiothreitol (DTT) 1mM, 1X Halt protease and phosphatase inhibitor cocktail) complemented with Triton X-100 0.1% for 5min on ice. After centrifugation at 1,500rcf, 5min, 4°C, the supernatant was clarified by high-speed centrifugation (18,000rcf, 4°C, 15min) to obtain the cytoplasmic fraction. The pellet was washed once with Buffer A and then incubated in Buffer B (EDTA 3.2mM, DTT 1mM, Halt protease and phosphatase inhibitor cocktail) for 30min on ice. After centrifugation (1,700rcf, 5min, 4°C), the supernatant was collected as the soluble nuclear fraction. The pellet (chromatin fraction) was washed once with Buffer B and resuspended in the same buffer. The whole cell extracts and chromatin-enriched fractions were then sonicated (10 pulses of 1s at 40% amplitude with a Sonics Vibra Cell Ultrasonic processor) and Laemmli buffer was added in order to have a final protein concentration of 2µg/µL and 0.5µg/µL respectively. The detection of pChk1 (S345, Cell signalling 2341, Rabbit), Chk1 (Santa Cruz sc 8408, Mouse), Actinin (MBL 05-384, Mouse), MCM2 (Abcam ab-4461, Rabbit), p-MCM2 S40 (Abcam ab133243, Rabbit), ORC2 (MBL M055-3, Mouse) Lamin A/C (Santa cruz sc-7293, Mouse) and Tubulin (Sigma T5168, Mouse) was done by running SDS-page gels, transferring on PVDF membranes, blocking with 5%milk, incubating with primary antibody (in TBS-T, adapted dilutions) followed by secondary antibody (MBL 70765,Mouse or MBL 70745 Rabbit, 1/10 000 in TBS-T) and finally revealing thanks to ECL (Biorad 170-5161) under the ChemiDoc imaging system (BioRad).

### Replication timing analysis

10-20 millions of exponentially growing mammalian cells (with DMSO or aphidicolin) were incubated with 0.5mM BrdU (Abcam, #142567), protected from light, at 37°C for 90 minutes. After washing in PBS, cells were fixed in 75% final cold EtOH and stored at -20°C. BrdU labeled cells were incubated with 80μg/mL Propidium Iodide (Invitrogen, P3566) and with 0,4 mg/ml RNaseA (Roche, 10109169001) for 15min at room temperature and 150 000 cells were sorted in early (S1) and late (S2) S phase fractions using a Fluorescence Activated Cell Sorting system (FACSAria Fusion, Becton Dickinson) in Lysis Buffer (50mM Tris pH=8, 10mM EDTA, 0.5% SDS, 300mM NaCl) and stored at -20°C until the following steps. DNA from S1 and S2 fractions of sorted cells was extracted using Proteinase K treatment (200µg/ml, Thermo Scientific, EO0491) followed by phenol-chloroform extraction and sonicated to a size of 500-1,000 base pair (bp), as previously described (39). Immunoprecipitation was performed using IP star robot at 4°C (indirect 200µl method, SX-8G IP-Star® Compact Automated System, Diagenode) with an anti-BrdU antibody (10μg, purified mouse Anti-BrdU, BD Biosciences, #347580). Denatured DNA was incubated for 5 hours with anti-BrdU antibodies in IP buffer (10mM Tris pH=8, 1mM EDTA, 150mM NaCl, 0.5% Triton X-100, 7mM NaOH) followed by an incubation for 5 hours with Dynabeads Protein G (Invitrogen, 10004D). Beads were then washed with Wash Buffer (20mM Tris pH=8, 2mM EDTA, 250mM NaCl, 1% Triton X-100). Reversion was performed at 37°C for 2 hours with a solution containing 1% SDS and 0.5mg Proteinase K followed, after the beads removal, by an incubation at 65°C for 6 hours in the same solution. Immunoprecipitated BrdU labeled DNA fragments were extracted with phenol-chloroform and precipitated with cold ethanol. Control quantitative PCRs (qPCRs) were performed using oligonucleotides specific of mitochondrial DNA, early (BMP1 gene) or late (DPPA2 gene) replicating regions (10, 39). Whole genome amplification was performed using SeqPlex™ Enhanced DNA Amplification kit as described by the manufacturer (Sigma-Aldrich, SEQXE). Amplified DNA was purified using PCR purification product kit as described by the manufacturer (Macherey-Nagel, 740609.50). DNA amount was measured using a Nanodrop. Quantitative PCRs using the oligonucleotides described above were performed to check whether the ratio between early and late replication regions was still maintained after amplification. Early and late nascent DNA fractions were labelled with Cy3-ULS and Cy5-ULS, respectively, using the ULS arrayCGH labeling Kit (Kreatech, EA-005). Same amounts of early and late-labeled DNA were loaded on human DNA microarrays (SurePrint G3 Human CGH arrays, Agilent Technologies, G4449A). Hybridization was performed as previously described (39). The following day, microarrays were scanned using an Agilent C-scanner with Feature Extraction 9.1 software (Agilent technologies). To determine the replication domains and do the comparative analysis in different conditions, the online platform specific for replication timing data START-R (40) was used, with biological duplicates for each condition. The output bed file gave the list of significantly impacted genomic regions (ADVANCED or DELAYED) and the report of number and percentage of genomic regions impacted. We also used the output replication timing smooth files to identify the early (RT > 1) the mid (−1 < RT < 1) and late (RT < -1) replicating regions.

### BrdU ChIP-qPCR

The same protocol as for replication timing was performed until BrdU immunoprecipitation (IP). For the BrdU-IP, 140μL of IP buffer (Tris pH8 50mM, EDTA 2mM, NaCl 300mM, Triton 1%, H2O qsp, 14mM NaOH extemporaneously) and 10µg of the monoclonal anti-BrdU antibody (BD Biosciences, 347580) were added to DNA and incubated on rotating wheel overnight at 4°C. 1.5mg of magnetic beads (Dynabeads™ Protein G, Thermofisher 1004D) previously washed with PBS (15min on wheel at RT) and IP buffer was added to the mix and incubated on wheel 2 hours at 4°C. After washing twice with 800μL of Buffer B (Tris pH8 20mM, EDTA 2mM, NaCl 250mM, Triton 0.2%, H2O qsp) and with 800μL of Tris pH8 10mM, beads were resuspended in 100μL of Tris pH8 10mM. Immuno-precipitated DNA was then recovered by a reversion step with a solution containing 1% SDS and 0.5mg Proteinase K for 2h at 37°C while shaking. The supernatant was incubated overnight at 65°C while shaking. A final phenol-chloroform purification was performed and DNA concentration was measured with Nanodrop technology before performing qPCR. qPCR were performed for the specific amplification of 2 early and 2 late replicating control regions, 3 ADV aRTIL and amplicon from neo-synthesized mitochondrial DNA for normalization (**Table S3**). StepOne technology was used to do the qPCR. For each genomic region amplified, we quantified the percentage of S1 and S2 after normalization with mitochondrial DNA (41).

### Gene expression microarrays

Exponentially growing cells (with DMSO or aphidicolin) were harvested and RNAs were extracted with RNeasy plus mini kit (Qiagen). RNAs quality and quantity were controlled using Nanodrop ND-1000 and Bioanalyzer 2100 Expert from Agilent. cDNAs were prepared according to the standard ThermoFisher protocol from 100ng total RNA (GeneChip™ WT PLUS Reagent Kit Manual Target Preparation for GeneChip™ Whole Transcript (WT) Expression Arrays User Guide). Following fragmentation, 5.5 µg of single stranded cDNA were hybridized on Human Clariom S Arrays in GeneChip Hybridization Oven 645 for 16 hr at 45°C. The arrays were washed and stained in the Affymetrix Fluidics Station 450. Arrays were scanned using the GeneChip Scanner GC3000 7G and images were analysed using Command Console software to obtain the raw data (values of fluorescent intensity). The data were analysed with TAC (Transcriptome Analysis Console, version 4.0.2.15) from ThermoFisher. Microarrays were normalized with the “Robust Multichip Analysis” (SST-RMA) method. Statistical analysis allowed tagging of genes according to the fold change (FC) and the p-value adjusted together with ANOVA approach.

### ATAC-seq

100,000 exponentially growing RKO cells were trypsinized, washed in PBS and treated with 1:100 volume of RNase-free DNase (QIAGEN) and DMEM media for 30min at 37°C in the incubator. Cells were trypsynized, washed in PBS and resuspended in 500μL of ice-cold cryopreservation solution (50% FBS, 40% DMEM, 10% DMSO), transferred into a 2mL cryotubes and frozen in a pre-chilled Mr. Frosty container at -80°C overnight or more before sending to Active Motif to perform ATAC-seq assay. The cells were then thawed in a 37°C water bath, pelleted, washed with cold PBS, and tagmented as previously described (42), with some modifications based on (43). Briefly, cell pellets were resuspended in lysis buffer, pelleted, and tagmented using the enzyme and buffer provided in the Nextera Library Prep Kit (Illumina). Tagmented DNA was then purified using the MinElute PCR purification kit (Qiagen), amplified with 10 cycles of PCR, and purified using Agencourt AMPure SPRI beads (Beckman Coulter). Resulting material was quantified using the KAPA Library Quantification Kit for Illumina platforms (KAPA Biosystems), and sequenced with PE42 sequencing on the NextSeq 500 sequencer (Illumina). Analysis of ATAC-seq data was very similar to the analysis of ChIP-Seq data. Reads were aligned using the BWA algorithm (mem mode; default settings). Duplicate reads were removed, only reads mapping as matched pairs and only uniquely mapped reads (mapping quality >= 1) were used for further analysis. Alignments were extended in silico at their 3’-ends to a length of 200 bp and assigned to 32-nt bins along the genome. The resulting histograms (genomic “signal maps”) were stored in bigWig files. Peaks were identified using the MACS 2.1.0 algorithm at a cutoff of p-value 1^e-7^, without control file, and with the –nomodel option. Peaks that were on the ENCODE blacklist of known false ChIP-Seq peaks were removed. Signal maps and peak locations were used as input data to Active Motifs proprietary analysis program, which creates Excel tables containing detailed information on sample comparison, peak metrics, peak locations and gene annotations. To annotate the ATAC-seq peak value and coverage within genomic regions of interest, we used Merge BedGraph and AnnotateBed bedtools functions respectively (Galaxy Version 2.29.2). We then normalized the three biological replicates values across all genomic regions (Early, Mid, Late, ADV and DEL) by a 2way ANOVA Sidak’s multiple comparisons test.

### Processing ChIP-seq data from public databases

ChIP-seq, pDamID and RepOri data were download from ENCODE, GEO, 4DN project and Replication domain data base respectively (**Table S4**). The epigenetic marks coverage of each given regions list (Early, Mid, Late, ADV, DEL) was calculated using AnnotatedBed bedtools function (Galaxy Version 2.29.2). The mean for each epigenetic mark coverage in given regions was calculated to generate clustering tree heatmap based on Pearson correlations with ClustVis software (44).

### Fluorescent DNA halo

400,000 cells were harvested after synchronization and treated with nuclei buffer (10 mM Tris at pH 8, 3 mM MgCl2, 0.1 M NaCl, 0.3 M sucrose, protease inhibitors) plus 0.5% Nonidet P40 for 5-10 min on ice (depending on cell line). Nuclei were attached to coverslips using cytospin (1500-1800 rpm for 5-10 min, depending on cell line); stained with DAPI (2 mg/mL for 4 min); and immersed in a buffer containing 25 mM Tris (pH 8), 0.5 M NaCl, 0.2 mM MgCl2, 1 mM PMSF, and protease inhibitors for 1 min, then in Halo Buffer (10 mM Tris at pH 8, 2 M NaCl, 10 mM ethylene diamine tetra acetic acid [EDTA], 1 mM DTT, protease inhibitors) for 4 min. After two washing steps with wash buffer 1 (25 mM Tris (pH 8), 0.2 M NaCl, and 0.2 mM MgCl2) for 1 min, and with buffer 2 (buffer 1 without NaCl) for 1 min extracted nuclei were fixed in 2% formaldehyde for 10 min and processed for immunofluorescence. Images containing about 200 halo per condition acquired with a Nikon Ni-E microscope and a DS-Qi2 camera with 64X objective and MFHR (Maximum Fluorescence halo Radius) was measured in Image J software.

## RESULTS

### Low replication stress differentially impacts cancer and non-tumour cells

In order to evaluate cellular responses to mild replication stress, we treated cells with 0.2µM aphidicolin and DMSO as a control. We used 6 well characterized cell lines that are common models (HCT116, RKO, U2OS, K562, MRC5-N and RPE-1) and differ in tissue origin, tumorigenicity, differentiation stage and molecular characteristics such as oncogene expression, genetic instability type and telomere maintenance mechanisms (**Table S1**). The duration of aphidicolin treatment was adapted to each cell line in order to treat a maximum number of cells in S phase for a single generation. We selected the longest treatment duration ensuring that no cells were treated during two consecutive cell cycles (**Table S2** and **Figure S1a**). This optimal treatment duration was also determined in order to analyse the effect of replication stress in the daughter cells released from the drug.

Analysis of the S phase checkpoint induction at the end of aphidicolin treatment, by monitoring Chk1 phosphorylation on serine 345, revealed a low level of checkpoint activation compared to acute replication stress (HU, 2mM, 2h), with nevertheless a tendency towards higher p-Chk1 level in cancerous cells (**Figure S1b,c**). Furthermore, we noticed a global accumulation of cells in the S phase under aphidicolin treatment, which again was more pronounced in cancerous cells (**Figure S1d,e**).

To study RT under low replication stress, BrdU was added to the culture medium before cell sorting into Early (S1) and Late S-phase (S2) fractions, and the neo-synthesized DNA was hybridized on human whole genome microarrays, as previously described (39, 45, 46) (**Figure S2a**). RT differential analyses were performed on biological replicate experiments using the START-R suite software (40) and only significant modifications between aphidicolin and control condition were retained.

It has been reported that the RT profile of somatic cells is closely related to the cell type and tissue origin (10). By performing hierarchical clustering of RT for the different cell lines, we found that, in absence of replication stress, non-tumour cells clustered together (cluster 1) and are separated from cancer cells (cluster 2) (**Figure 1a**). In cluster 2, we noticed that the replication timing of RKO is closer to HCT116, consistent with the fact that they are both colon cancer cell lines with microsatellite instability due to mismatch repair deficiency (MMR-) **(Table S1)**. Upon aphidicolin treatment, the two distinct clusters of cancer and non-tumour cells remain apart but distances measuring relatedness between cancer cells are altered. Indeed, the RT of RKO cells appears to be closer to that of K562 cells and that of HCT116 to be closer to U2OS (**Figure 1b**). Overall, this observation indicates firstly that without any replication stress, RT itself can discriminate non-tumour from cancer cells, and secondly, that aphidicolin treatment affects differentially the identity of cancer cells since we observed that the RT of the two colon cancer cell lines (RKO and HCT116) are now further apart among the cancer cells cluster 2 (**Figure 1b**).

**Figure 1:**
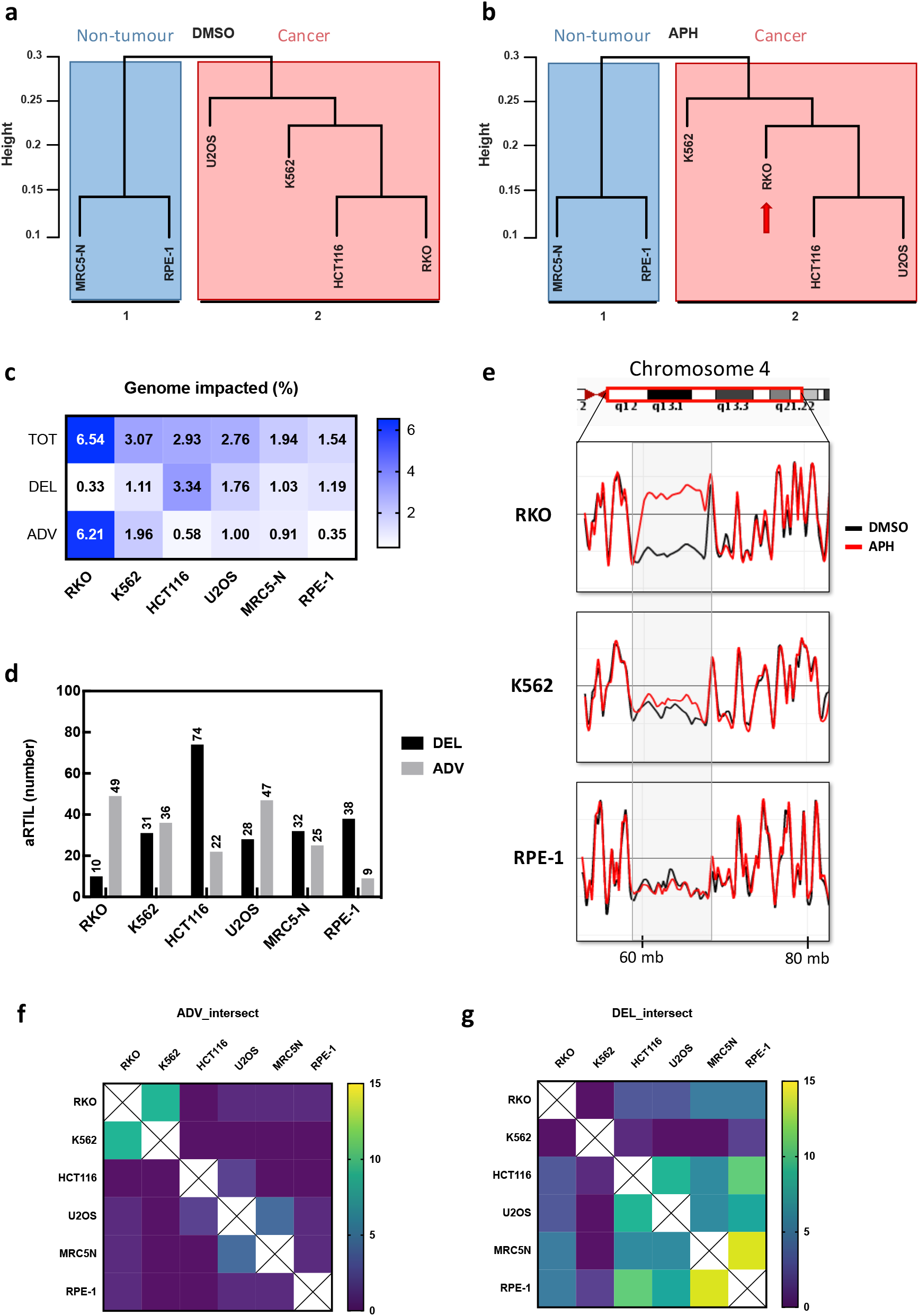
Low replication stress differentially impacts cancer and non-tumour cells. **a** Cluster dendogram based on p-values reflecting RT signatures in DMSO condition of the six cell lines. Distance: correlation and clustering method: average. **b** Cluster dendogram based on p-values reflecting RT signatures in APH condition of the six cell lines. Distance: correlation and clustering method: average. **c** Heatmap representing the coverage (in % of the genome) of impacted genomic regions: total, delayed or advanced (TOT, DEL and ADV) for each cell line. **d** Quantification of the number of genomic regions significantly impacted (DEL, black and ADV in grey) by aphidicolin treatment. **e** Screenshot of Loess-smooth RT profiles for the same region in chromosome 4 for RKO, K562 and RPE-1 cells. The dark lines correspond to replication timing of control (DMSO) replication timing (2 independent replicates) and the red lines are replication timing of APH treated cells (2 independent replicates). **f** Heatmap with intersections (the sum of the number of common domains) for ADV genomic regions between cell lines. **g** Heatmap with intersections (the sum of the number of common domains) for DEL genomic regions between cell lines.

### Advance RT signature in cancer cells

We recapitulated the genome wide percentage of altered RT in the different cell lines in response to aphidicolin treatment (**Figure 1c**). Normal RPE-1 and MRC5-N cells were the least impacted cell lines with 1.54% and 1.94% of the genome undergoing RT alterations respectively (**Figure 1c**) while the RT in cancerous cells was globally more impacted by low replication stress. We noticed that the RKO cell line showed the highest response with 6.54% of the genome impacted (**Figure 1c,e** and large visualization of chromosomes in **Figure S3**).

Analysis of the aphidicolin-RT-impacted loci (aRTIL) led to the identification of regions with significant RT delays (DEL aRTIL), as previously reported (30, 32) (**Figure 1c,d**). In RPE-1 and HCT116, these DEL aRTIL represent the majority of impacted loci with 38/47 and 74/96 loci respectively. Importantly, in all cell lines, we also observed significant RT switches towards earlier RT (RT advances; ADV aRTIL) (**Figure 1c-e**). These ADV aRTIL represent the main type of RT changes in RKO cells, with the largest domain coverage (**Figure 2c** and **Figure S3**) and strongest amplitude (**Figure 1e** and **Figure S3**). By performing BrdU-ChiP-qPCR on three independent domains in the RKO cell line, we confirmed that these newly-identified advanced domains are effectively replicated earlier (relative to BrdU incorporation) upon aphidicolin treatment (**Figure S2b**).

**Figure 2:**
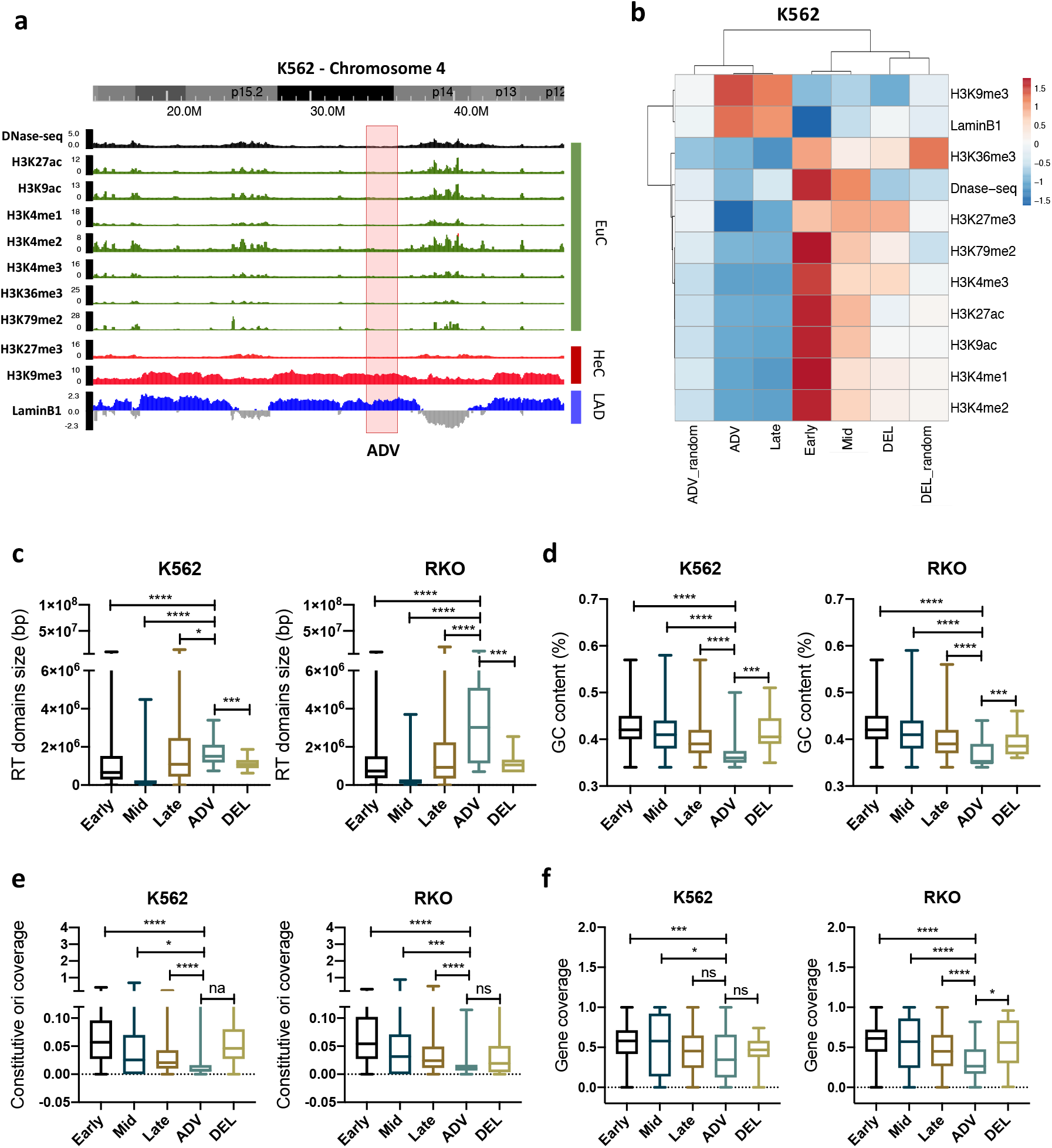
ADV aRTIL occur in heterochromatin late replicated chromosomic regions. **a** Screenshot of WashUEpigenome Browser for chromosome 4 in K562 with an example of ADV aRTIL (in red). **b** Heatmap and clustering trees based on Pearson correlations for epigenetic marks coverage in K562 regions (ClusVis Software, ENCODE ChiP-seq data). **c** Boxplots of RT domains size (in kb) for Early, Mid, Late, ADV aRTIL and DEL aRTIL in K562 and RKO cell lines. **d** Boxplots of GC content in Early, Mid, Late, ADV aRTIL and DEL aRTIL genomic regions in K562 and RKO cell lines. **e** Boxplots of constitutive origins coverage in Early, Mid, Late, ADV aRTIL and DEL aRTIL genomic regions in K562 and RKO cell lines. **f** Boxplots of gene coverage in Early, Mid, Late, ADV aRTIL and DEL aRTIL genomic regions in K562 and HCT116 cell lines. Statistics (for all the boxplots): Wilcoxon rank sum test ***p<0.005, *p<0.05, ns when p>0.05.

To explore the potential similarities of RT modifications between cell lines, we quantified the number of common DEL and ADV aRTIL (**Figure 1f,g**). We found 9 common large ADV domains in RKO and K562 cells and 4 common ADV aRTIL in U2OS and MRC5-N. The few ADV domains detected in RPE-1 and HCT116 were mainly specific to each cell line. The two non-tumour cell lines (RPE-1 and MRC5-N) shared a majority of DEL aRTIL (14 common) and also had many in common with U2OS (8 and 6 with RPE-1 and MRC5-N respectively) and HCT116 cell lines (11 and 6 with RPE-1 and MRC5-N respectively). In contrast, DEL aRTIL found in RKO and K652 cells were poorly shared with other cell lines.

Collectively, these data demonstrate that RT is significantly modified in cancer cells for a subset of genomic domains in response to mild replication stress. RT of non-tumour cells RPE-1 and MRC5-N was less affected, together with the lower induction of the S-phase checkpoint. Importantly, these data reveal for the first time that, in addition to RT delays, replication stress can also induce RT advances. Finally, the overlapping of aRTIL between cell lines highlights two main distinct RT modification signatures: the first one in RKO and K562 cell lines, characterized by major shared RT advances in specific genomic domains, and the second for U2OS, HCT116, MRC5-N and RPE-1 that are sharing similar RT delayed genomic regions.

### ADV aRTIL occur in heterochromatin late replicated chromosomal regions

To investigate whether aRTIL are associated to specific epigenomic signatures, we next analysed histone modifications and other epigenomic features such as Lamina-Associates-Domains (LADs) and chromatin accessibility (DNase-seq). Given that the most advanced domains in RKO are shared with K562, we used public epigenomic data on K562 to characterize ADV aRTIL. First, this approach allowed us to validate our experimental RT through the expected enrichment of the typical chromatin marks of Early, Mid and Late replicated regions (**Figure 2a,b**). For instance, Early replicated regions are enriched in euchromatin marks such as H3K27ac and H3K9ac, while Late replicated regions correlate with heterochromatin marks (H3K9me3) and with LaminB1 (LADs) (**Figure 2a,b**). We also confirmed that chromatin accessibility (DNAse-seq) decreases from Early to Late replicated regions. Interestingly, chromatin features of the ADV aRTIL are similar to those of Late replicated domains and we noticed that these are even more enriched in H3K9me3 and in LaminB1, while being poorer in H3K27me3, and display very low chromatin accessibility (**Figure 2a,b**). This result suggests that ADV aRTIL domains belong to constitutive heterochromatin while DEL aRTIL are more likely Mid-replicated regions (**Figure 2b**).

We next investigated if genomic features can help distinguish ADV from DEL aRTIL. In accordance with the epigenomic features, we showed that in untreated cells, the large majority of regions converted to ADV aRTIL are replicated in Late S-phase while those changed to DEL aRTIL are mainly replicated in the Early/Mid S-phase (**Figure S4a**). At the genomic sequence level, we observed that ADV aRTIL are large regions (**Figure 2c**) that share similar features with Late replicated regions, such as poor GC content (**Figure 2d**), few constitutive origins (**Figure 2e**) and low gene abundance (**Figure 2f**). In contrast, DEL aRTIL are enriched in GC content, origins and gene coverage that characterize Early and/or Mid replicated regions (**Figure 2d-f** and **Figure S4c-e**).

### The ADV aRTIL signature is related to CFS but is not targeted by DNA damage signalling

Given that CFS are the most sensitive chromosomal regions to replication stress and that RT delays have been described in these fragile loci, we wondered if aRTIL would also overlap with CFS. To answer this question, we analysed the overlap between aRTIL and the CFS already identified (30). We found a clear enrichment of CFS within aRTIL in all 6-cell lines, with 22-44% of CFS being located within aRTIL (**Figure 3a-c**). The percentage of RT delays within CFS was higher in the HCT116, MRC5-N and RPE-1 cell lines (**Figure 3b**). Quite surprisingly, we also identified ADV aRTIL associated to CFS, notably for the cancerous RKO and K562 cells. Therefore, besides the well-documented delayed replication dynamics within CFS, replicative stress can also induce RT advances in these particular regions.

**Figure 3:**
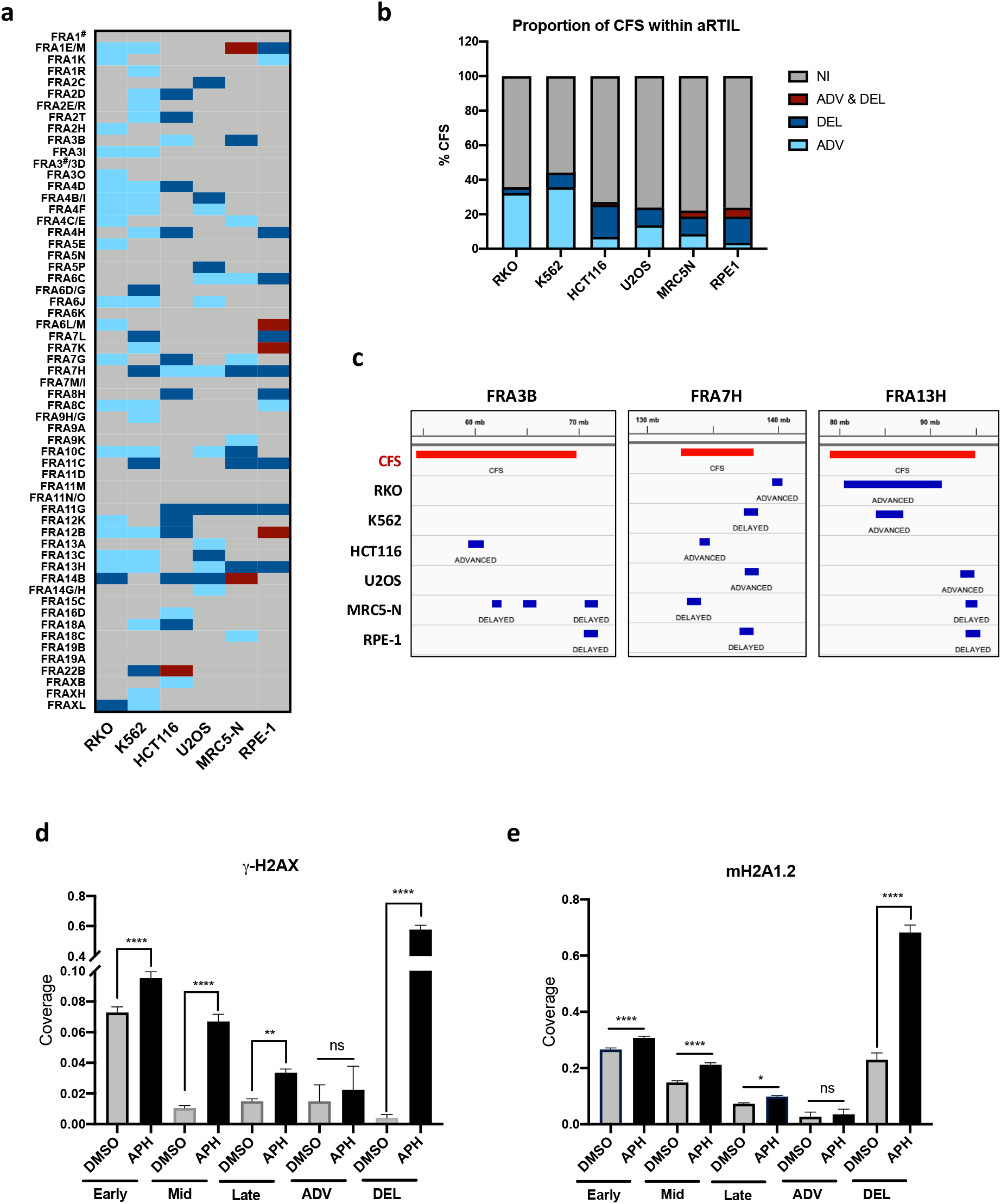
The ADV aRTIL signature is closely related to CFS but is not targeted by DNA damage signalling. **a** Categorical heatmap to visualize impacted CFS by aRTIL in all cell lines: ADV & DEL in dark red, DEL in dark blue, ADV in light blue and non-impacted (NI) in grey. **b** Histogram representing the proportion (in %) of CFS within aRTIL. ADV & DEL in dark red, DEL in dark blue, ADV in light blue and non-impacted (NI) in grey. **c** Genome Browser (IGV) snapshots to visualize aRTIL (in blue) in all cell lines for three CFS (in red): FRA3B, FRA7H and FR13H. **d** Histogram representing the coverage of γ-H2AX histone mark (ChIP-seq data) on Early, Mid, Late, ADV aRTIL and DEL aRTIL genomic regions in DMSO- and APH-treated conditions. Statistics: Two-way ANOVA, Multiple comparison: ****p<0.0001, **p<0.01, *p<0.05, ns p>0.05. **e** Histogram representing the coverage of mH2A1.2 histone mark (ChiP-seq data) on Early, Mid, Late, ADV and DEL aRTIL genomic regions in DMSO- and APH-treated conditions. Statistics: Two-way ANOVA, Multiple comparison: ****p<0.0001, **p<0.01, *p<0.05, ns p>0.05.

It has been recently reported that macroH2A1.2, a variant from the canonical H2A (47), having roles both in replication stress response and in cell fate decisions (48–51), is more abundant at CFS than non-fragile regions of the genome. In response to mild aphidicolin treatment (0.5 µM), its enrichment is directly correlated with γ-H2AX peak coverage (52). Using ChIP-seq data from this study, we analysed the coverage of histone variants γ-H2AX and mH2A1.2 in K562 with or without aphidicolin treatment. As we did for epigenomic marks, we compared ADV and DEL aRTIL coverage with Early, Mid and Late S-phase regions. We confirm that, without aphidicolin, Early-S regions are the most enriched in γ-H2AX and mH2A1.2 histone variants (**Figure 3d-e)** (53). As expected, aphidicolin induces a significant increase in γ-H2AX and mH2A1.2 in all control regions. Interestingly, aphidicolin treatment does not modify the coverage of γ-H2AX and mH2A1.2 within ADV aRTIL whereas it induces a strong increase of these two histone modifications coverage within DEL aRTIL (**Figure 3d-e**). This suggests that, while DEL regions are likely prone to DNA damage under replication stress, ADV aRTIL could be protected from DNA damage or more efficiently repaired.

Taken together, these results show that replication stress differently affect aRTIL. In contrast to DEL aRTIL, ADV aRTIL are associated to a low level of DNA damage response (DDR) signalling.

### Low replication stress impacts the regulation of genes involved in chromatin organization

It has been established that RT switches in human cells can be linked to developmental genes expression (3, 13, 54, 55). Nonetheless, the exact correlation between RT and gene expression is not entirely clear. Indeed, several studies have discovered genomic sequences that do not fit the general correlation between gene expression and RT (56–58). Therefore, we checked if, in our experimental condition, aphidicolin has a global impact on gene expression and if this could be related to RT modifications. We performed gene-expression profiling by microarray in RKO cells using the same conditions as for RT analysis. We found that aphidicolin treatment has a mild impact on gene expression, with 14 genes significantly differentially expressed (APH DOWN or APH UP genes) (**Figure 4a and Figure S5a**). We analysed APH DOWN genes by performing a Gene Ontology (GO) enrichment analysis and found these genes were enriched in chromatin and nucleosome organization, gene silencing and cellular differentiation pathways (**Figure S5b**). We did not observe particular enrichment in GO pathways for APH UP genes (FDR > 0.01). Nevertheless, we noticed an up-regulation of the transcription factor gene ZBTB38 which is a biomarker for prostate cancer (59) and that plays an important role in the regulation of DNA replication, cell cycle and cell fate (60, 61). Importantly, the expression of genes that fell inside aRTIL was not affected by low replication stress (**Figure 4b**). Altogether, we concluded that while RT modifications under low replication stress are not related to modification of gene expression within aRTIL, replication stress induces a specific down-regulation of genes involved in chromatin organization.

**Figure 4:**
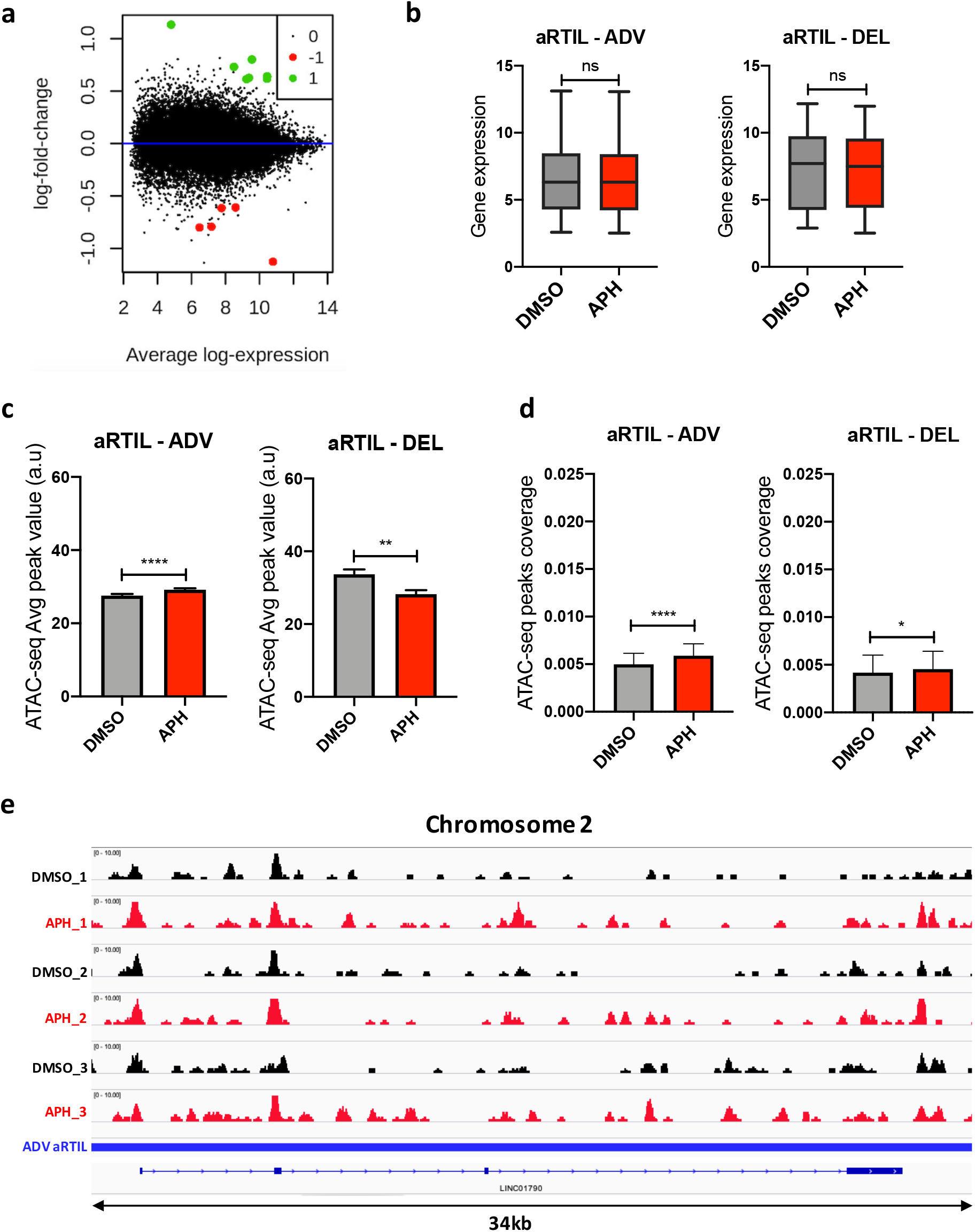
Aphidicolin modulates chromatin accessibility within ADV aRTIL without impacting gene expression. **a** Scatterplot for RNA-ChIP data in RKO cells representing significantly UP (green) and DOWN (red) -regulated genes in response to aphidicolin. **b** Boxplots measuring the expression of genes within ADV (left) and DEL (right) aRTIL in DMSO (grey) and APH (red) conditions. Statistics: Wilcoxon matched-pairs signed rank test ns p>0.05. **c** Comparison of ATAC-seq peak value within ADV and DEL aRTIL regions between DMSO (grey) and APH (red) conditions. **d** Comparison of ATAC-seq peak coverage within ADV and DEL aRTIL between DMSO (grey) and APH (red) conditions. Statistics (N=3): Wilcoxon matched-pairs signed rank test ****p<0.0001, **p<0.01, *p<0.05. **e** Screenshot of integrative genome viewer (IGV) session with ATAC-seq triplicates bigwig files on chromosome 2 within a specific ADV aRTIL (194,728,595-194,763,435kb, LINC01790).

### Aphidicolin modulates chromatin accessibility within ADV aRTIL

Since we identified APH DOWN genes in RKO cells involved in chromatin and nucleosome organization, we examined the impact of aphidicolin on chromatin structure by performing the Assay for Transposase Accessible Chromatin with high-throughput sequencing (ATAC-seq), a method for assaying chromatin accessibility genome-wide (62, 63). We observed a global remodelling of chromatin accessibility under aphidicolin treatment demonstrated by a lower strength of ATAC-seq peaks and an increase ATAC-seq peaks coverage in both Early and Late replicated genomic regions (**Figure S5c,f**). More importantly, we observed a significant increase of both ATAC-seq peaks strength and coverage within ADV aRTIL while this was not the case within DEL aRTIL (**Figure 4c-e** and **Figure S5e**). Thus we can conclude that under low replication stress chromatin accessibility is specifically increased within ADV aRTIL.

We also investigated the chromatin accessibility of the UP-regulated genes under aphidicoline treatment. In contrast to ADV aRTIL, we did not find an increase of ATAC-seq peaks value within these specific genes (**Figure S5g**). This result further supports that differential gene expression is not associated to chromatin remodelling and RT modifications induced by aphidicolin treatment.

Overall, this approach enabled us to demonstrate that aphidicolin treatment induces a local increase of chromatin accessibility within ADV aRTIL that can be linked to RT modifications while also inducing whole genome chromatin remodelling.

### ADV aRTIL can be transmitted to daughter cells

RT is faithfully established at the beginning of the G1 phase in each cell cycle, at a precise time named the “timing decision point” or TDP (64, 65). In G1, RT setting up is dependent on 3D nuclear replication domains organisation through chromatin loop formation mediated by the Rif1 protein (66). Moreover, at the G1/S transition, chromatin loops are also maintained by the transient recruitment of pre-replication complex proteins and active origins to the nuclear matrix (NM) (67–70).

We first wondered if RT changes under replication stress in mother cells can be preserved beyond the G1 phase and thus transmitted to daughter cells. To answer this question, we released the 6 cell lines from aphidicolin or DMSO treatment for the appropriate duration (**Table S2**) and analysed the RT of daughter cells in S phase (N+1). Our results indicate that DEL and ADV aRTIL observed in mother cells were no longer detected in the next cell generation of K562, HCT116, U2OS, MRC5-N and RPE1 (**Figure 5a and Figure S6a**). Strikingly, in RKO cells, the majority (28 of 49) of the strongest and largest ADV aRTIL were transmitted to the next cell generation and we noticed that the amplitude of these RT advances was less pronounced in daughter cells (**Figure 5a,b** and **Figure S3**). Conversely, DEL aRTIL returned to normal RT in RKO daughter cells, comparable to untreated cells. These results indicate that while the majority of the RT modifications within aRTIL are reversible and tend to be eliminated through the G1 phase, severe ADV aRTIL can be transmitted to the next generation.

**Figure 5:**
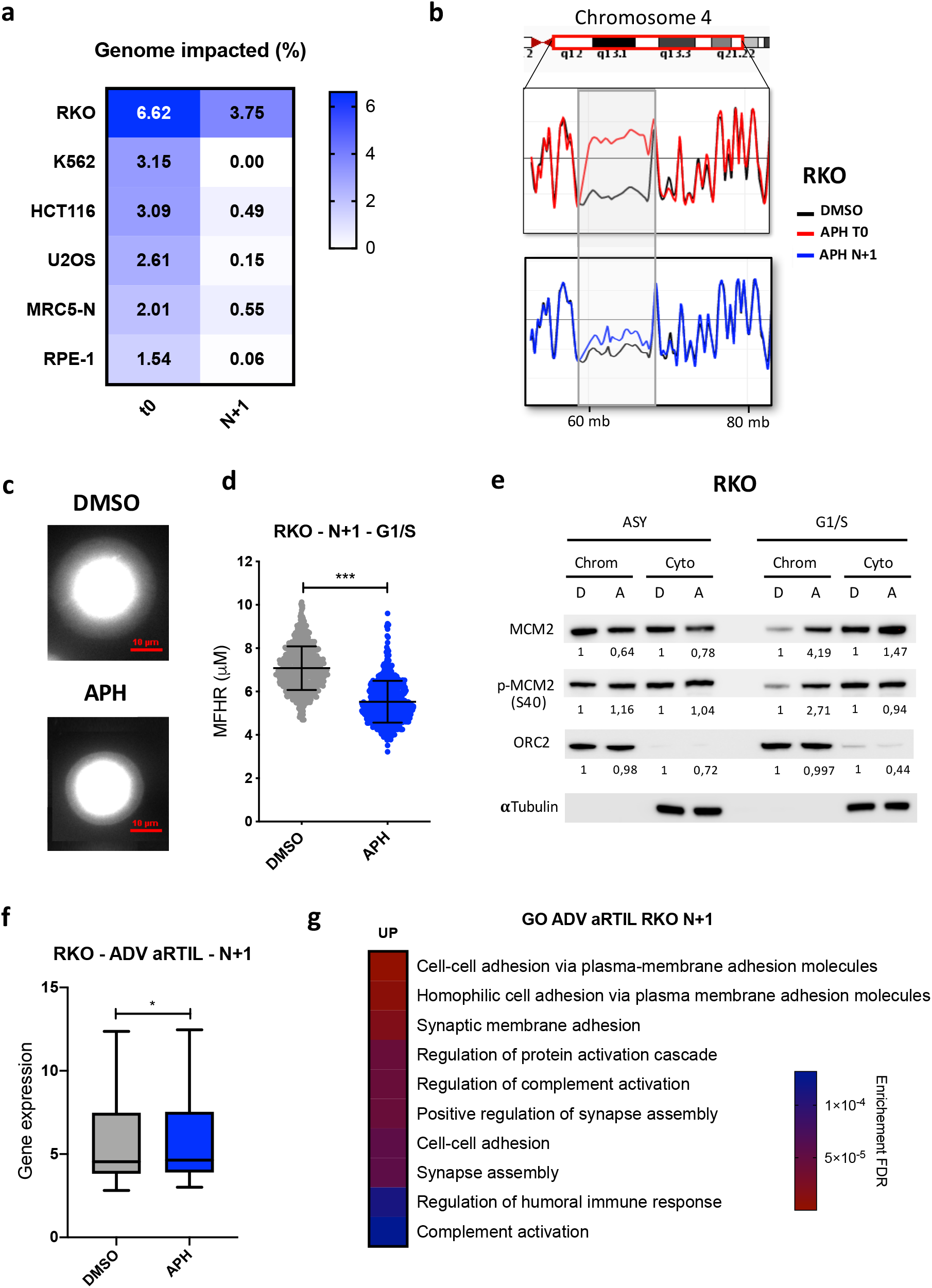
ADV aRTIL can be transmitted to daughter cells. **a** Heatmap representing the coverage (in %) of impacted genomic regions in mother cells (t0) or released daughter cells (N+1) for the six cell lines. **b** Screenshots of Loess-smooth replication timing profiles for the same region in chromosome 4 for RKO, in mother (APH T0) and daughter cells (APH N+1). The dark lines correspond to replication timing of control (DMSO) replication timing, the red line is replication timing of T0 APH-treated cells and the blue line is the replication timing of N+1 daughter cells released from APH treatment. **c** Visualization and **d** Quantification of DNA Halo size (MFHR) in RKO G1/S synchronized daughter cells released from DMSO or aphidicolin treatment. Statistics (N=3): Unpaired t test with Welch’s correction ***p<0.001. **e** Western blot on chromatin (Chrom) and cytoplasmic (Cyto) protein fractions to quantify the amount of MCM2, p-MCM2 and ORC2 in asynchronous cells (ASY) and cells synchronized in G1/S with L-mimosine. The fold change between DMSO and APH condition measured in three independent experiments is reported in the figure. **f** Boxplot measuring the expression of genes within ADV aRTIL in DMSO and APH released daughter cells (N+1). Statistics: Wilcoxon matched-pairs signed rank test *p<0.05. **g** Gene ontology for RKO N+1 ADV aRTIL UP-regulated genes (ShinyGO, P-val cutoff: FDR < 0.05).a

To investigate if the persistence of ADV aRTIL in RKO daughter cells could be linked to changes in chromatin loop organisation in G1 phase, we performed a fluorescent DNA halo experiment to evaluate the chromatin loops size in G1/S. We used RKO cells as positive control and RPE-1 cells as negative control. We measured the maximum fluorescence halo radius (MFHR) formed around the nuclear matrix (NM) and noticed a significant shrinkage of DNA loops in the aphidicolin-released RKO cells (**Figure 5c-d**) while no effect was measured in RPE-1 cells (**Figure S6 b-c**). To test if this reduced halo size correlates to an increase in licensed origins, we quantified the loading of pre-replication complex components onto the chromatin under the same conditions. Our results clearly show an increase of MCM2 and p-MCM2 loading onto the chromatin in G1/S of RKO aphidicolin-released daughter cells, while this was not the case in RPE-1 cells (**Figure S6d**), further supporting the results from the DNA halo experiment (**Figure 5e**). The transmission of ADV aRTIL in RKO daughter cells is therefore associated to a decrease in chromatin loop size and an increase in pre-RC proteins loading in G1 phase, predicting greater activation of replication origin in the next S phase.

Finally, we wondered if chromatin remodelling during the G1 phase and RT advances in RKO daughter cells would modulate gene expression. Even though the expression of genes within ADV aRTIL was not significantly impacted in mother cells, we noticed an up-regulation of genes within ADV aRTIL in daughter cells released from replication stress (**Figure 5f**). Therefore, we performed a gene ontology analysis to determine if these ADV aRTIL genes were involved in a specific molecular pathway(s) and found a strong enrichment for cell-to-cell adhesion and synapse assembly pathways (**Figure 5g**).

Together these data show a correlation between the transmission of RT advances and chromatin structure modification at the time of the G1 phase. In addition, we observed that persistence of RT advances in daughter cells is associated to an increase of gene expression within these specific genomic regions. Overall, we revealed that replication stress in mother cells directly affects chromatin accessibility and replication timing, which will further affect the cell fate of the next generation.

## DISCUSSION

DNA replication is tightly regulated in order to guarantee the accurate transmission of genomic information from mother to daughter cells. Challenging structures of DNA are very often encountered by the replication fork and oncogene expression or metabolism changes can also play a deleterious role in DNA replication program efficiency by inducing replication stress. Thus, regardless of the sources, replication stress will give rise to many cellular responses that will depend on the genetic background of the cell. Importantly, replication stress was identified at the early stage of cell transformation and cancer development (71, 72) and persists in cancer cells after the selection steps (73). As replication stress compromises genome stability, cancer cells have to adapt in order to survive and maintain their proliferation. Due to a high level of genomic instability, cancer cells can lose some of the classical mechanisms involved in DDR, for example through well-known mutations in BRCA1/2 repair genes. However, many compensatory processes take place in cancer cells to allow survival (74–76), also conferring resistance to chemotherapeutic treatments.

In the present work, we demonstrated that one of the components of the replication program, RT, is affected in response to low dose of aphidicolin. Besides the already described DNA replication delays following replication stress induced by the inhibition of replicative DNA polymerases α, δ and ε, we revealed here that aphidicolin also promotes RT advances (**Figure 1 and S2, S3**). In order to understand how the same replication stress induces such opposite effects, we characterized DEL and ADV aRTIL at both the genomic and epigenomic level. DEL aRTIL are normally replicated in the Early/Mid S-phase, they harbour many genes and a high number of constitutive origins of replication (**Figure S4**). These features explain the high probability for replication forks to be affected by aphidicolin treatment and hampered by transcription activity, leading to DNA damage signalling associated with high levels of γ-H2AX and macroH2A1.2 (**Figure 3d,e**).

ADV aRTIL are mainly observed in RKO and K562 cancer cells, even if they are also detected in the other cell lines. These regions correspond to Late replicated heterochromatin with few and poorly expressed genes, low number of constitutive origins and a low coverage of the two DDR histone marks γ-H2AX and mH2A1.2, not significantly increased under low replication stress (**Figure 2, 3**). Interestingly, in RKO cells, we demonstrated higher chromatin accessibility within ADV aRTIL in the mother cells that is not linked to higher gene expression (**Figure 4**). Altogether, ADV aRTIL appear to be heterochromatin flexible regions which might be resistant to replication stress.

Under low dose of aphidicolin, the decrease in DNA replication fork velocity can be compensated by the firing of dormant origins, preventing under-replication (77–82). Interestingly, it was demonstrated that in cancer cells replication origin usage is more flexible (83). Moreover, cancer cells rely on a higher rate of origin licensing protein expression (84, 85), allowing a more efficient usage of dormant origins in response to replication stress (86). Taking our data into consideration, we can imagine that, in cancer cells, the higher chromatin accessibility described within ADV aRTIL favours the access of DNA replication and repair proteins to these specific heterochromatin regions, leading to activation of dormant origins and earlier DNA replication. It was described that activation of replication stress-induced dormant origins within a given S phase can be persistent in the next S phase (78). This result is consistent with our data in RKO cells, in which we still observe RT advances in daughter cells after aphidicolin release accompanied by a reduction in DNA halo size and an increase in MCMs loading to chromatin at the G1/S transition (**Figure 5**). Altogether, we propose that some cancer cells are able to modify their replication program and display a higher flexibility of chromatin organisation and replication origin usage in order to respond to replication stress. In addition, we observed that genes within ADV aRTIL are up-regulated in daughter cells (**Figure 5**). These results would suggest that the chronic replication stress described during the first step of cancer development could be involved in the acquisition of replication timing modifications that subsequently lead to changes in the transcription level of specific genes. Thus, low replication stress would change cell fate of cancer cells whose chromatin is more flexible.

RT modifications are historically associated with organism development and cellular differentiation (3, 4, 8, 87). Furthermore, the pluripotent capacity of mammalian cells has been linked to genome plasticity (88, 89) as well as deficiency in Lamin A protein expression (90, 91). In our study, we observed that replication stress induced higher level of RT modifications in cancer cells. Interestingly, we showed that RT of the cancerous RKO cell line is strongly affected by aphidicolin treatment, inducing major ADV aRTILs (**Figure 1c,d,e and S3**). We linked this particular phenotype to the poorly differentiated status of RKO cells together with a very low expression of Lamin A/C (**Table S1, Figure S7**). In agreement with this, we noticed that the K562 cell line, which had an aRTIL signature close to the RKO cells, is poorly differentiated and also expressed a low level of Lamin A/C (**Table S1, Figure S7**). Overall, we propose that the less differentiated cancer cells are, the more they will harbour RT advances in response to replication stress, as a sign of their more flexible chromatin organization.

Finally, we unveiled a new mechanism in response to replication stress that may have a strong impact on gene expression and cell identity, mainly for cancer cells. Importantly this work paves the way for future studies investigating the molecular effectors involved in advancing replication timing in cancer cells, with great potential as new targets to prevent cancer cells adaptation to replication stress leading to therapy resistance.

## ACCESSION NUMBERS

DNA replication timing, ATAC-seq and microarray gene expression data are available under accession numbers GSE156618, GSE156552 and GSE156521, respectively.

## SUPPLEMENTAL DATA

Supplementary Data are available at NAR online.

## ACKNOWLEDGMENT

We are grateful to the Technologic platform from CRCT (Manon Farcé for cytometry, Marie Tosolini and Juan-Pablo Cerapio-Arroyo for the bio-informatics analysis). The authors also thank Nathalie Marsaud (GeT-Biopuces, Toulouse Biochip Facility) for her help and advice about the experimental protocol of Microarray Transcriptomic. We thank Marie Jeanne Pillaire, Guillaume Labrousse, Gaëlle Legube, Chunlong Chen and Frédéric Bantignies for their helpful advice and discussions throughout this project. We also want to thank Sarah McClelland, Laure Crabbe, Jean-Yves Masson and Emmanuelle Bitoun for their precious feedback and proofreading.

