## Supplemental figures and tables for "Low replication stress leads to specific replication timing advances associated to chromatin remodelling in cancer cells"

### Supplemental data

| Cell line | RKO | HCT116 | U2OS | K562 | MRC5-N | RPE-1 |
|---|---|---|---|---|---|---|
| **Gender** | Female | Male | Female | Female | Male | Female |
| **Cancer type** | Carcinoma | Carcinoma | Osteosarcom a | Leukemia | Non-cancerous | Non-cancerous |
| **Tissue** | Colon | Colon | Bone | Bone marrow | Lung | Retina |
| **Differentiation** | Poorly differentiated | Poorly Unable to differentiate | Moderately differentiated | Undifferentiated progenitor | Embryonic | Terminally differentiated |
| **Morphology** | Epithelial | Epithelial | Epithelial | Hematopoietic | Fibroblast | Epithelial |
| **Molecular characteristics** | p53+ (WT)<br>MMR-<br>Telomerase<br>CIN- | p53+ (WT)<br>MMR-<br>Telomerase<br>CIN- | p53+ (WT)<br>MMR+<br>ALT<br>CIN+ | p53-<br>MMR+<br>Telomerase<br>CIN+ | p53+ (WT)<br>MMR+<br>Normal<br>CIN- | p53+ (WT)<br>MMR+<br>hTERT<br>CIN- |

**Table S1: Cellular and molecular characteristics of the 6 human cell lines used for the study.** RKO and HCT116, which are two colon cancer cell lines, are highlighted in orange, in green the U2OS cell lines, from osteosarcoma and in yellow the K562 cells lines from haematological cancer chronic myeloid leukaemia. The two human cells from healthy tissues MRC5-N and RPE-1 are coloured in grey (dark and light) and are respectively from lung and retina. Cellular background regarding differentiation state, morphology and finally some molecular characteristics like p53 status or telomere maintenance mechanism are listed in this table. MMR: Mismatch repair, CIN: chromosome instability.

| | **RKO** | **K562** | **HCT116** | **U2OS** | **MRC5-N** | **RPE-1** |
|---|---|---|---|---|---|---|
| **Duration of treatment (T0)** | 16h | 12h | 15h | 15h | 15h | 12h |
| **Duration of release (N+1)** | 13h | 18h | 14h | 15h | 12h | 13h |

**Table S2: Optimised durations of treatment with aphidicolin (T0) and release in a fresh media (N+1) for the 6 cell lines.** Experimentally, BrdU was added all along aphidicolin treatment, in order to follow S phase treated cells, and EdU was incorporated only at the end to detect S phase cells at this time point. For the T0, we selected the longest times of treatment that maintained a minimum of BrdU positive cells in G1 phase, indicating that no cell was treated twice (**Figure S1a**). For the N+1, cells were release in a fresh media for the indicated duration, in order to have a maximum of aphidicolin treated cells reaching the next S phase.

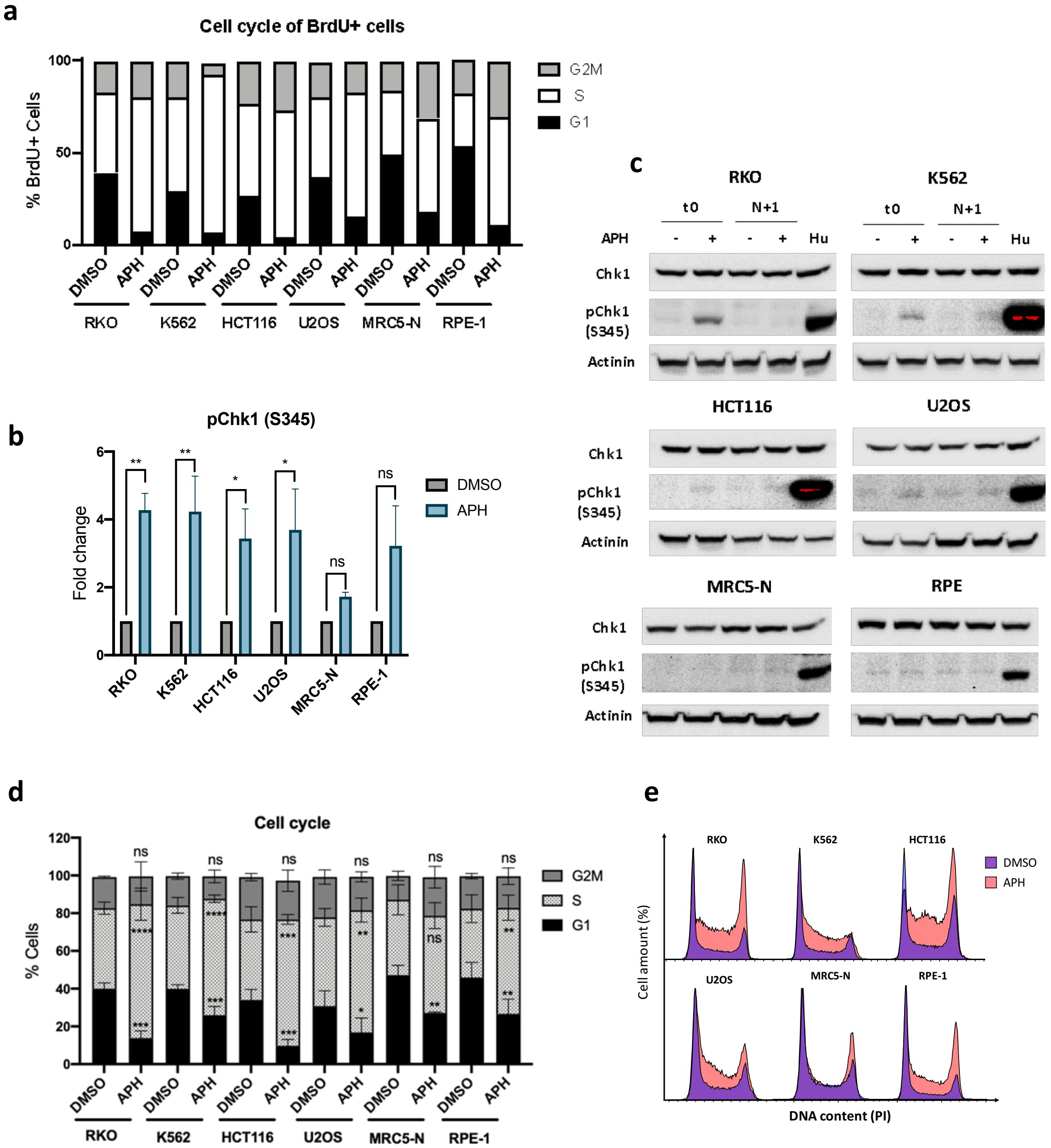


**Figure S1: Low replication stress differentially impacts cancer and non-tumour cells. a** Stacked histograms representing the percentage of BrdU positive cells in each cell cycle fraction (G1, S or G2M). Cell cycle analysis was assessed by FACS (DAPI, EdU and BrdU) labeling after BrdU incorporation during the treatment (indicated in **Table S2**) and 15min EdU pulse before cell harvesting and fixation. **b** Quantification of p-Chk1 intensity between DMSO and aphidicolin (APH) treatment detected by western blot. Fold change correspond to normalized APH pChk1 intensity (by Chk1 and Actinin intensities) divided by normalized DMSO pChk1 intensity. Statistics (N>4): Two-way ANOVA, Multiple comparison: **p<0.01, *p<0.05, ns when p>0.05. **c** Western blot of pChk1 (Serine 345), Chk1 and Actinin with and without APH for the first generation of cells (t0), daughter cells (N+1) or positive control Hu (2mM, 2h). **d** Stacked histograms representing cell cycle distribution (in percentage) between G1, S and G2/M phase. Statistics (N=3): Two-way ANOVA, Classical ****p<0.0001, ***p<0.005, **p<0.01, *p<0.05, ns when p>0.05. **e** Overlaps of DMSO (purple) and APH (pink) density plot showing cell cycle based on PI content by FACS analysis.

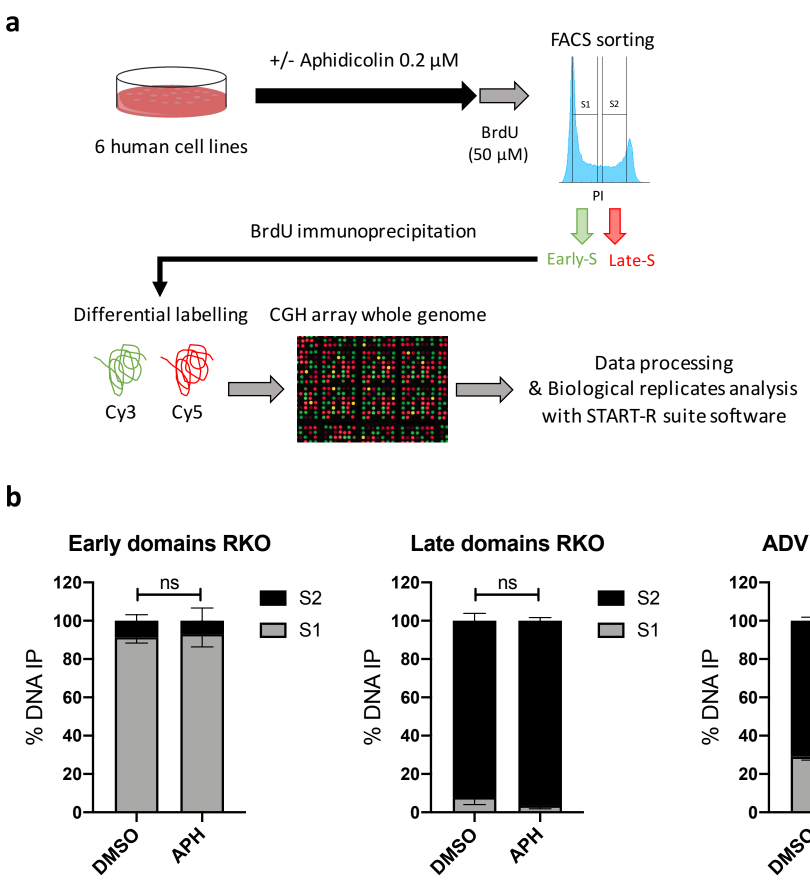


**Figure S2: Design and validation of replication timing experiment. a** Experimental protocol to assess whole genome-RT. Briefly, each cell line was treated with 0.2µM aphidicolin or DMSO for adapted duration and labelled with BrdU during 1h30 then FACS sorted into two S phase fractions (S1 for Early S and S2 for Late S). DNA from each fraction was immunoprecipitated with BrdU antibody, immunolabelled with Cy3 (S1) and Cy5 (S2), and then hybridized on whole human genome arrays. **b** BrdU-ChiP-qPCR after cell cycle sorting between Early (S1) and Late (S2) S phase. Stacked histograms represent the % of DNA IP for a given genomic domain in the S1 or S2 fraction. The first histogram recapitulates the results for the two Early control domains (E1 and E2), the second for the two Late control domains (L1, L2) and the last one for three advanced regions in the aphidicolin condition (ADV aRTIL). Statistics (three biological replicates, N=3): 2way ANOVA Sidak's multiple comparison test ****p<0.0001, ns when p>0.05.

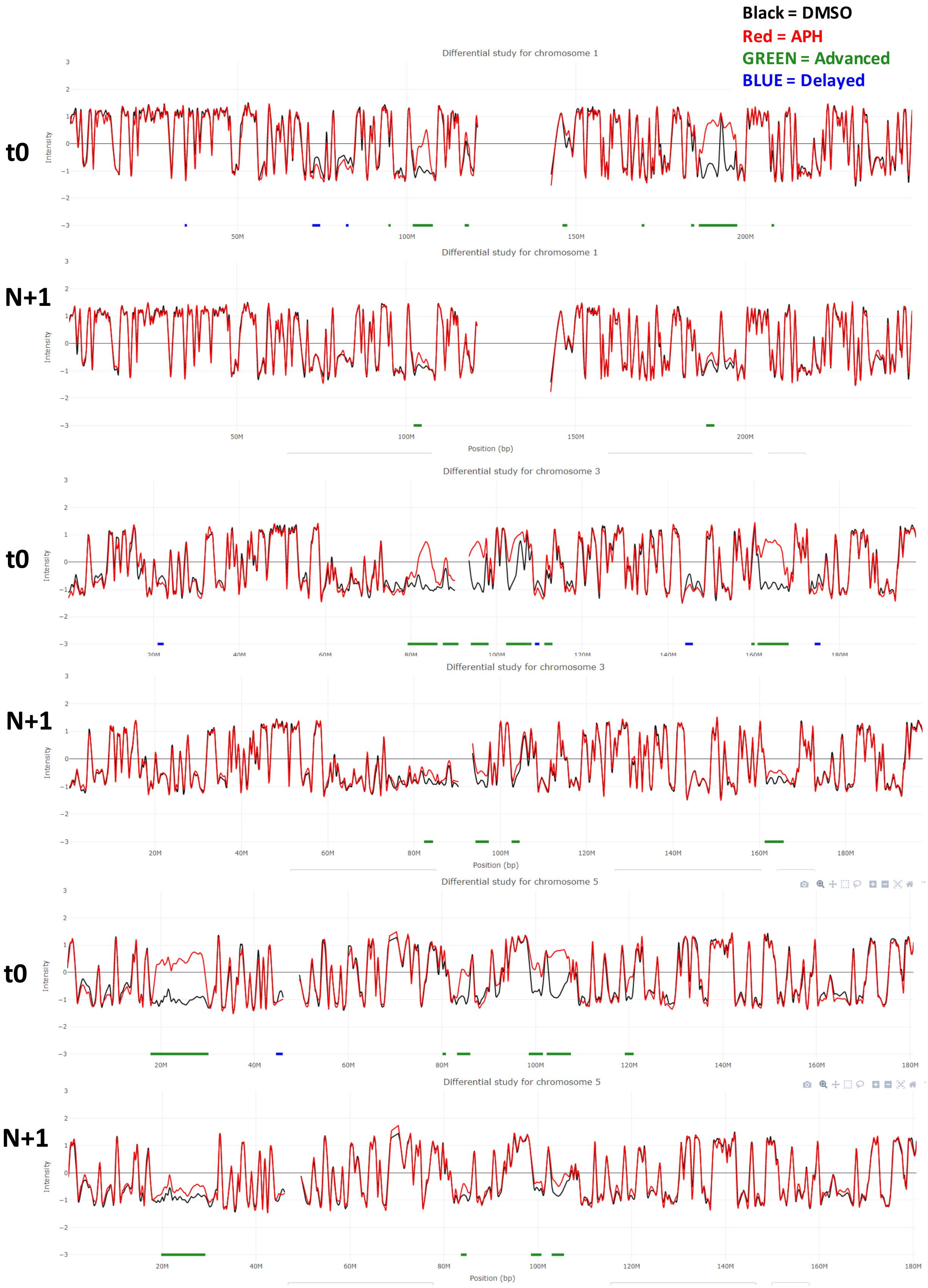
Black = DMSO
Red = APH
GREEN = Advanced
BLUE = Delayed
Differential study for chromosome 1
t0
N+1
Differential study for chromosome 3
t0
N+1
Differential study for chromosome 5
t0
N+1
Intensity
Position (bp)

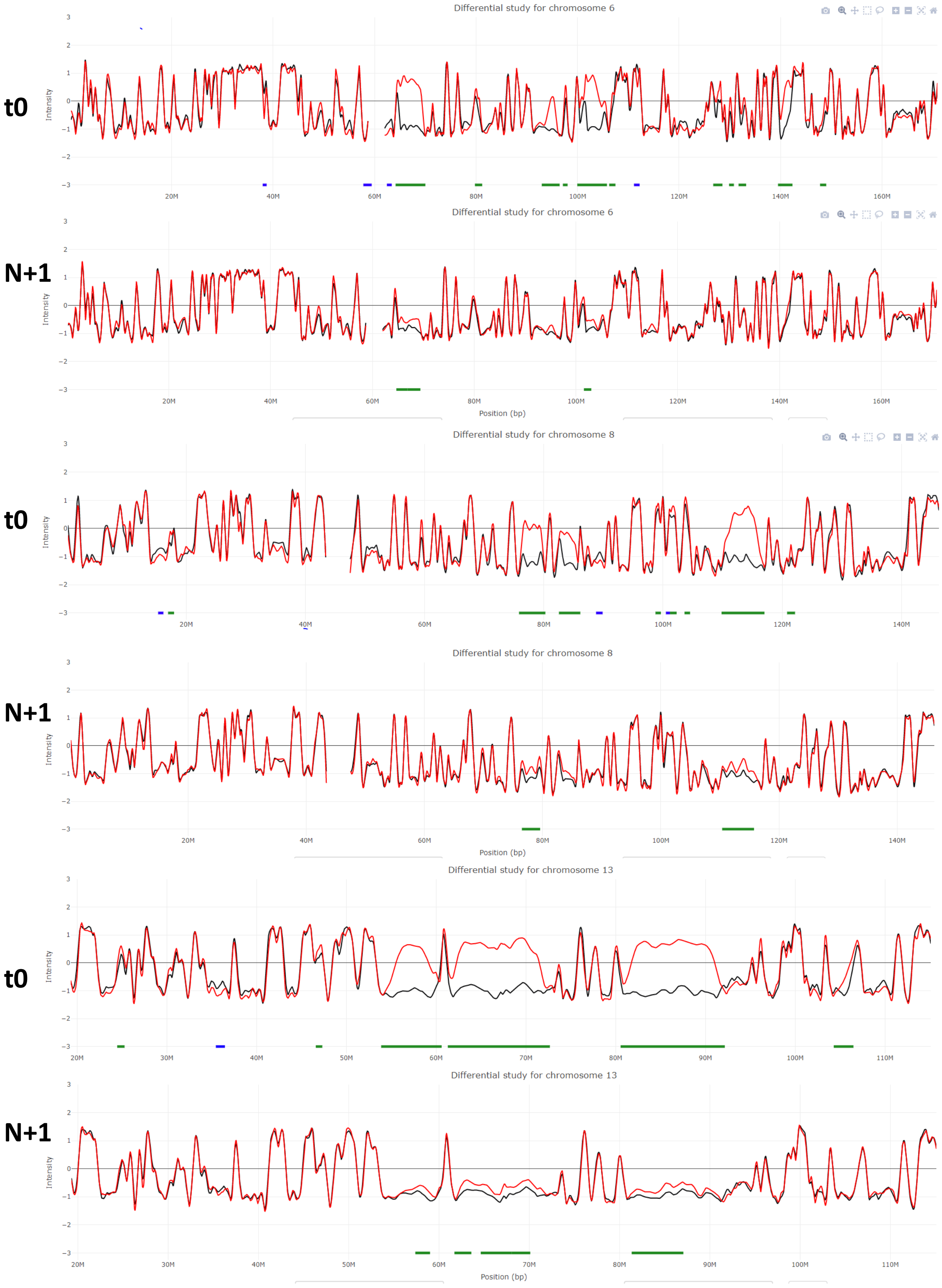


**Figure S3: DNA replication timing visualisation in RKO cells.** Screenshots of the 6 first chromosome visualisation of whole Loess-smooth RT profiles in mother (t0) and daughter cells (N+1). The dark lines correspond to RT of control (DMSO) replication timing and the red lines to RT of APH treated cells (2 independent replicates). ADV aRTIL are underlined in green and DEL aRTIL in blue.

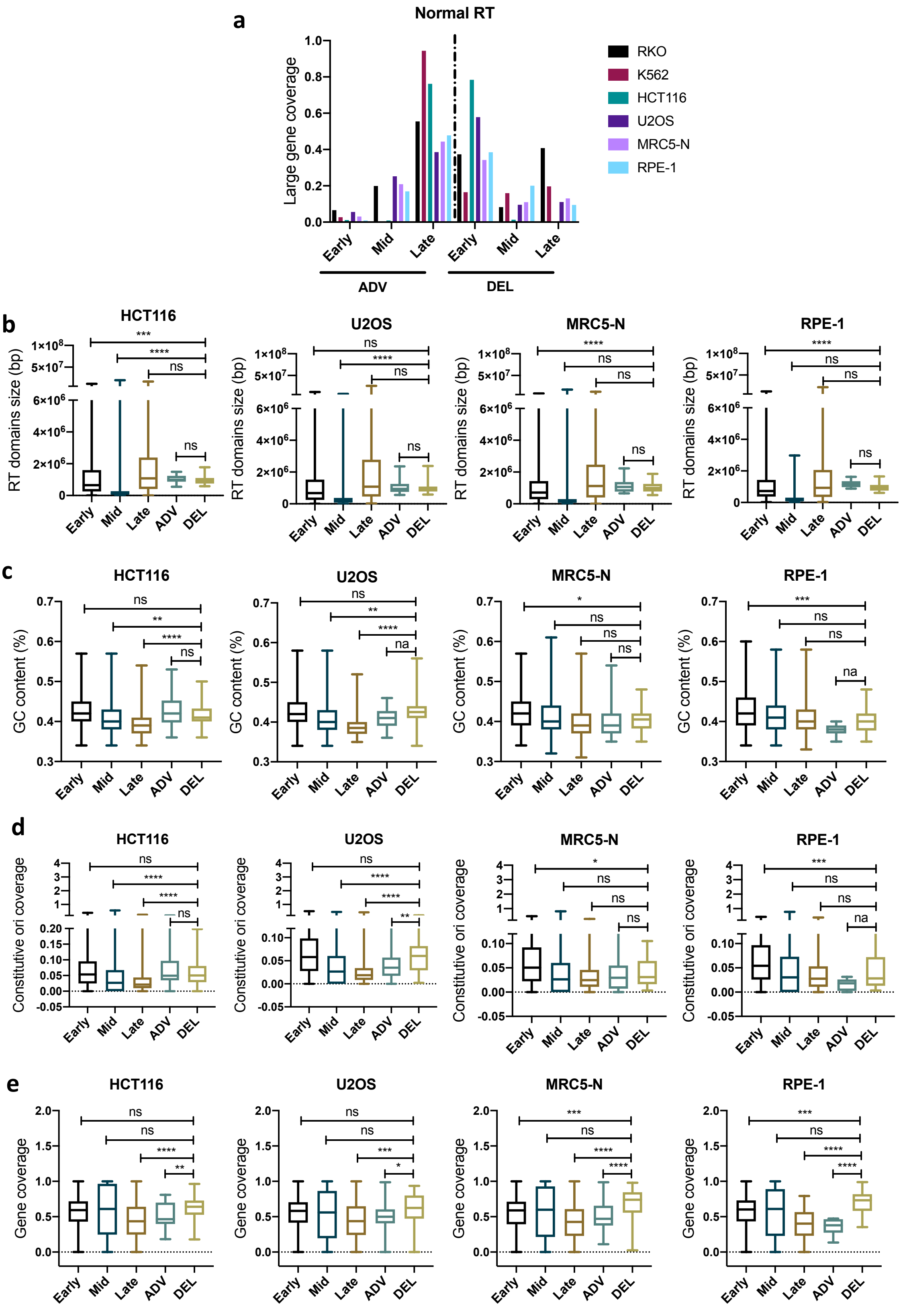


**<u>Figure S4</u>: aRTiL genomic characterisation. a** Histograms representing the normal replication timing (DMSO condition) of ADV and DEL aRTIL genomic regions in RKO, K562, HCT116, U2OS, MRC5-N and RPE-1. **b** Boxplots of RT domains size of Early, Mid, Late, ADV aRTIL and DEL aRTIL genomic regions in HCT116, U2OS, MRC5-N and RPE-1. **c** Boxplots of GC content (in percentage) in Early, Mid, Late, ADV aRTIL and DEL aRTIL genomic regions in HCT116, U2OS, MRC5-N and RPE-1. **d** Boxplots of constitutive replication origins coverage in Early, Mid, Late, ADV aRTIL and DEL aRTIL genomic regions in RKO, U2OS, MRC5-N and RPE-1. Statistics: Wilcoxon rank sum test ****p<0.0001, ns when p>0.05. **e** Boxplots of gene coverage in Early, Mid, Late, ADV aRTIL and DEL aRTIL genomic regions in HCT116, U2OS, MRC5-N and RPE-1. Statistics (for all the boxplots): Wilcoxon rank sum test ***p<0.005, *p<0.05, ns when p>0.05.

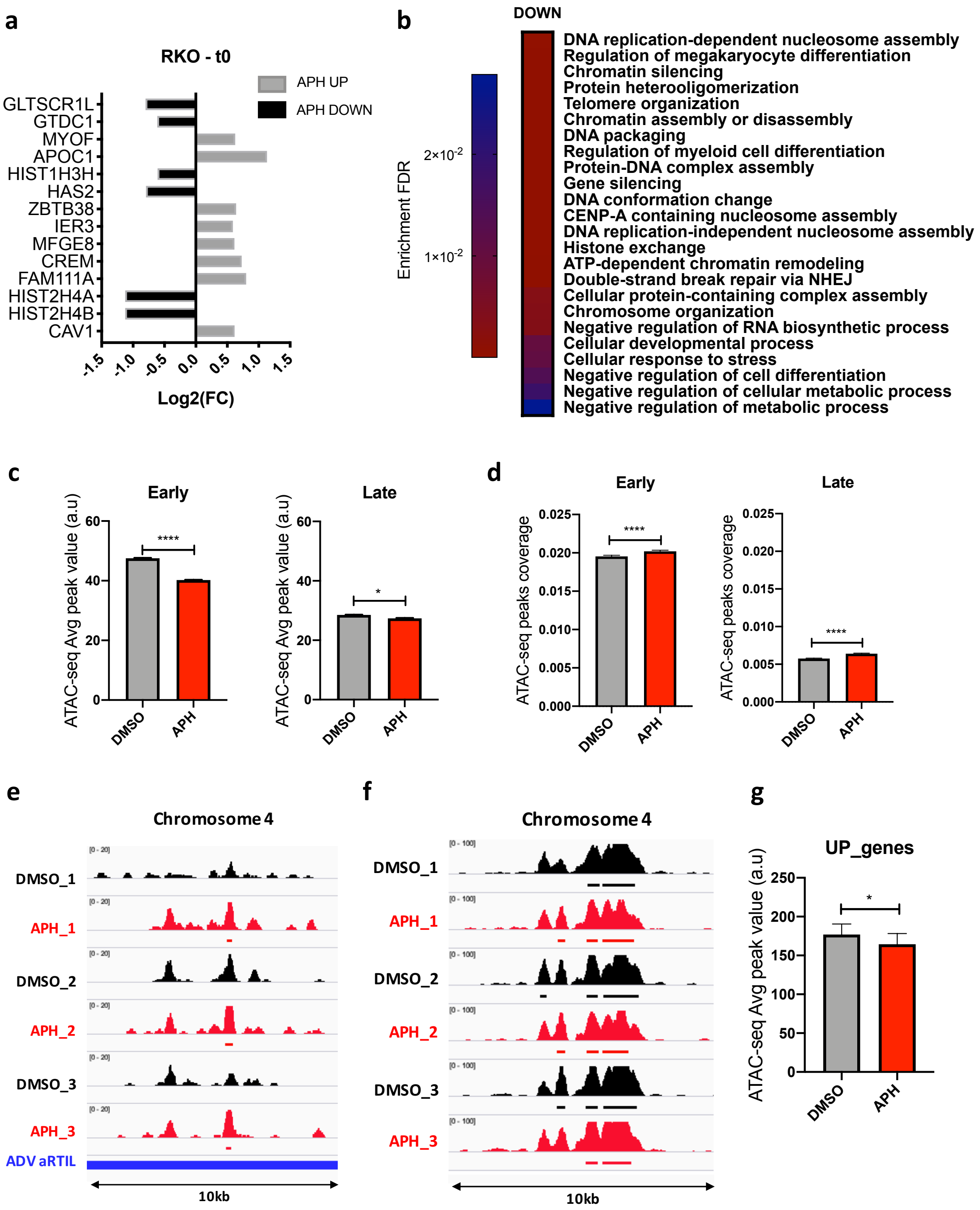


**Figure S5: Low replication stress impacts the regulation of genes involved in chromatin organization and modulates chromatin accessibility within ADV aRTIL.** **a** Names and values of log2 Fold change gene expression induced by aphidicolin for the most significantly impacted genes (Log2(FC) <0, APH DOWN, in black; Log2(FC) >0, APH UP, in grey). Statistics: ANOVA test p<0.05. **b** Gene ontology for DOWN-regulated genes (ShinyGO, P-val cutoff: FDR < 0.05). **c** Comparison of ATAC-seq peak value within Early and Late regions between DMSO (grey) and APH (red) conditions. **d** Comparison of ATAC-seq peak coverage within Early and Late replicating regions between DMSO (grey) and APH (red) conditions. Statistics (N=3): Wilcoxon matched-pairs signed rank test ****p<0.0001, *p<0.05. **e** Screenshot of integrative genome viewer (IGV) session with ATAC-seq triplicates bigwig files on chromosome 4 within a specific ADV aRTIL (91,820-91,830kb). **f** Screenshot of integrative genome viewer (IGV) session with ATAC-seq triplicates bigwig files on chromosome 4 within a random non impacted region (151,010-151,020kb). **g** Comparison of ATAC-seq Average peaks value (Avg peak value) between DMSO (black) and APH (grey) conditions within APH UP-regulated genes (UP-genes). Statistics (N=3): Wilcoxon matched-pairs signed rank test *p<0.05.

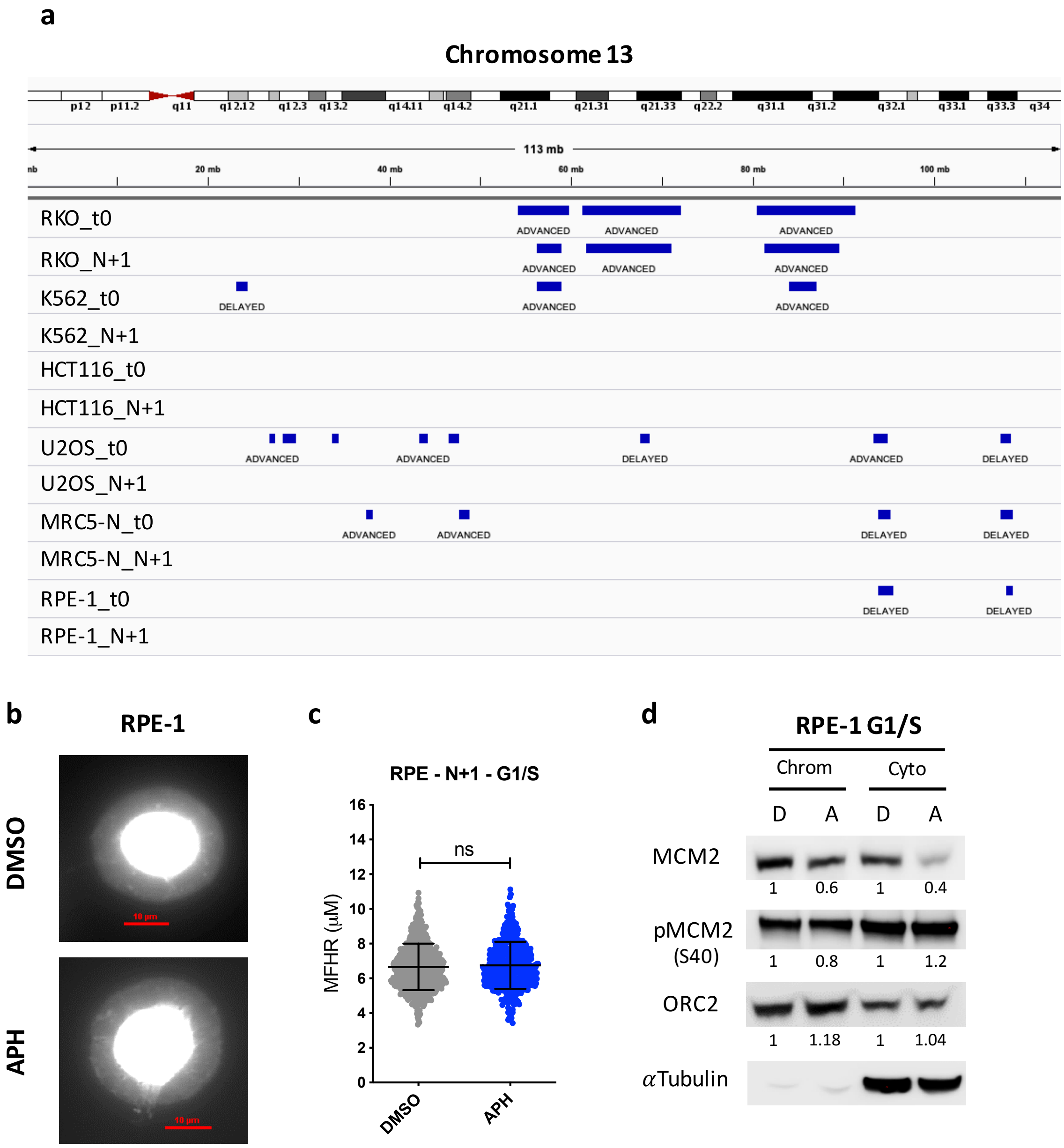


**Figure S6: ADV aRTIL can be transmitted to daughter cells. a** Screenshot of integrative genome viewer (IGV) session with aRTIL on chromosome 13 in mother (t0) and released daughter cells (N+1) for the six cell lines. **b** Visualization and **c** Quantification of DNA Halo size (MFHR) in RPE-1 G1/S synchronized daughter cells released from DMSO or aphidicolin treatment. Statistics (N=3): Unpaired t test with Welch's correction ns when p<0.05. **e** Western blot on chromatin (Chrom) and cytoplasmic (Cyto) protein fractions to quantify the amount of MCM2, p-MCM2 and ORC2 in the G1/S cells (synchronized with L-mimosine). The fold change between DMSO and APH conditions is reported on the figure.

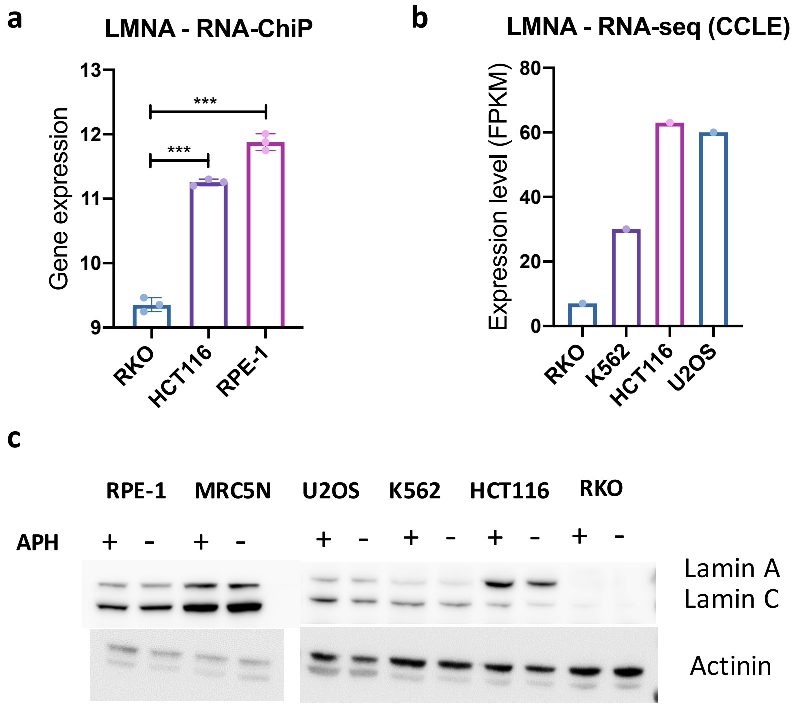


**<u>Figure S7:</u> Gene expression and protein level of LaminA/C in cancer and non-tumour cells. a** RNA-ChIP expression of LMNA gene in RKO, HCT116 and RPE-1. Statistics: Two-way ANOVA Tukey's multiple comparisons test ****p<0.0001. **b** Expression level (FPKM) of LMNA gene expression in RKO, K562, HCT116 and U2OS (RNA-seq values from the CCLE data set). **c** Western Blot for LaminA/C and actinin (for loading control) protein in the six cell lines aphidicolin treated (APH+) or not (APH-).

| Name | Forward sequence | Reverse sequence | chr | Start | End |
|---|---|---|---|---|---|
| L1 | CCCCATCCCCAGTTCTTTCC | TGGTGGGACTTGTGCTGTTT | chr17 | 67077058 | 67077088 |
| L1bis | TGGCACGTTCTTGTCACACT | AAGGTCCTCAGCCATTCAGC | chr17 | 67129038 | 67129068 |
| L2 | TCACTGCCAGTTCGACACAG | ACCCAGTCCCACATCACTCT | chr16 | 75360480 | 75360510 |
| L2bis | TGTGTCCATGCTGTGCCTAG | TGATGGAAGCAGCTACGTGG | chr16 | 75255289 | 75255319 |
| E1 | TGACTTCCGCTTCGAACCTC | GATGCTTGCACTCCCTCTGT | chr1 | 45058256 | 45058286 |
| E2 | TCCATCCTCAGGTCCTCGAG | AATGGCACGGTTCTCAGGAG | chr12 | 119955736 | 119955766 |
| ADV1 | GTTTCCTGGATGTTGCGCAG | CCAACAGAGACCCAGCAGAG | chr13 | 88356053 | 88356083 |
| ADV1bis | CAGCTTCACAGACCTCTCCG | GCTTGCATCACTGGAGTCGA | chr13 | 81884982 | 81885012 |
| ADV2 | GCCAAGCTCGCACAATGTTT | TCTTTCCACTTGCACCTGGG | chr8 | 114462446 | 114462476 |
| ADV2bis | GTTGCCCCATCACGAAAACC | GCATGACCCTGTATCTGCCC | chr8 | 114378227 | 114378257 |
| ADV3 | ATGGATTAGGCCCCGGTACT | CTCACTGCAATCCTGACGGT | chr1 | 187080568 | 187080598 |
| ADVbis | GAAGCTTCAGAGGCCGACAT | GGAGACCATCAATGCCCGTT | chr1 | 187034792 | 187034822 |
| Mito | CCTAGGAATCACCTCCCATTCC | GTGTTTAAGGGGTTGGCTAGGG | / | / | / |

**<u>Table S3:</u> Name, genomic sequences and location of BrdU-ChIP-qPCR probes.**

| | K562 |
|---|---|
| H3K4me1 | ENCFF320NKZ |
| H3K27ac | ENCFF770UZZ |
| H3K36me3 | ENCFF676RW |
| H3K27me3 | ENCFF369ZKL |
| H3K9ac | ENCFF842GQO |
| H3K4me3 | ENCFF729NQN |
| H3K9me3 | GSM607494 |
| H3K79me2 | ENCFF520XBR |
| H3K4me2 | ENCFF298QOP |
| DNase-seq | ENCFF433TIR |
| LaminB1 | 4DNFIHGCLSYW |
| gH2AX_NT | GSM2808044 |
| gH2AX_APH | GSM2808042 |
| mH2A1.2_NT | GSM2808015 |
| mH2A1.2_APH | GSM2808019 |

**<u>Table S4:</u> Reference of public ChIP-seq data used in this study.** Data from the ENCODE project are highlighted in yellow, those from the GEO dataset in blue and those from the 4DN project in green.
